# Complex carbohydrates catabolism capacity of bladder microbiota inhabiting healthy and overactive female bladders

**DOI:** 10.1101/2025.04.02.646827

**Authors:** Jean-Philippe F. Gourdine, Itallia V. Pacentine, Alec Barstad, Erin M. Dahl, W. Tom Gregory, Alan J. Wolfe, Tatyana A. Sysoeva, Lisa Karstens

## Abstract

Overactive bladder syndrome (OAB) is a poorly understood symptom complex that affects 40% of females over the age of 40, with clinical features including urinary urgency and incontinence. In addition to inflammation, oxidative stress, nerve damage and reduced blood flow, alterations in the urinary microbiome (urobiome), specifically in bladder bacterial diversity, have been reported to be associated with OAB. Bladder bacteria are members of the urobiome along with viruses, archaea, fungi, and protozoans. The urobiome metabolism, particularly in relationship to host complex sugars (glycans), has been investigated recently in terms of glycosaminoglycan (GAG) utilization. Nevertheless, other urinary free oligosaccharides (FOS) have not yet been explored in both OAB and urobiome contexts. Similarly, a comprehensive search of microbial genes involved in host glycan metabolism in the bladder of adult females with or without OAB has not yet been reported. In this study, we investigated urinary FOS by mass spectrometry in women without OAB (asymptomatic controls), with OAB without incontinence (dry OAB), or with OAB with incontinence (wet OAB or urgency urinary incontinence, UUI). We also questioned the ability of commensal bladder bacteria to digest these FOS and other glycans, using bioinformatic tools to query publicly available bladder genomes isolated from affected and unaffected adult females to identify genes that encode polysaccharide lyases (PL) and glycoside hydrolases (GH). Our results show that FOS are present in a similar level in affected and unaffected controls with a few exceptions: ten FOS were found to differ between the OAB dry groups and either the control (four) or UUI (six) groups. Our results indicate that bladder microbiota from adult females both with and without OAB have the genetic capacity to digest host glycans and dietary sugars with subtle differences. Bladder bacteria isolated from females with OAB possess more GH/PL genes for host mucins, whereas bladder bacteria from controls possess more GH/PL genes for GAG digestion. In the control group, specifically, the genus *Streptococcus* possessed genes for the PL8 and GH88 enzymes, known to be involved in host GAG digestion. These novel bioinformatic data can enable future biochemical exploration of the urobiome’s metabolism toward specific host glycans, such as GAGs, mucins O-glycans and N-glycans.

## Introduction

Overactive bladder syndrome (OAB) is a common symptom complex that negatively impacts the health and quality of life of over 16.5% American adults^1^. OAB is defined by a combination of lower urinary tract symptoms that include frequency and urgency of urination that can be either accompanied by incontinence or not (the so-called ‘wet OAB’ subtype [also known as urgency urinary incontinence, UUI] or ‘dry OAB,’ respectively). The etiology of OAB is not well understood, likely due to its putative multifactorial nature. In addition to inflammation, oxidative stress, nerve damage, and reduced blood flow, changes in the urinary microbiome (urobiome) composition have been associated with OAB^2–7^. Since the discovery of the urobiome, several studies have focused on urobiome comparison in adult females with OAB compared to adult females without lower urinary tract symptoms^3,8–11^. These studies found microbiome composition alterations and certain bacterial taxa associated with bladder symptom severity, response to treatment, or predictive of post-surgery urinary tract infection (UTI) risk but no clear marker of OAB has yet been identified^12^.

One underexplored aspect of the urobiome is the metabolic capacity of these resident urinary microbes. In contrast, the bladder environment itself is well understood, consisting of a bladder surface lining of known composition and a lumen containing bulk urine (Fig. 1). The urinary bladder lining is a unique protective barrier consisting of tightly bound umbrella urothelial cells covered with a protective mucosal layer composed of glycosylated proteins (proteoglycans, N-linked and O-linked glycoproteins), glycolipids, and other complex polysaccharides (glycosaminoglycans [GAGs]), collectively known as the glycocalyx^13,14^. The umbrella cells themselves are covered with uroplakins and mucins, glycoproteins that are heavily decorated primarily with N-glycans^15–17^ or O-glycans^18^, respectively. Mucins are either secreted or cell-associated O-glycosylated proteins that protect epithelial surfaces^18^, with MUC1 and MUC4 commonly found in the urothelium^19^ (Fig. 1). In related urinary conditions that combine urgency and frequency of urination with pain (Interstitial Cystitis [IC] and Bladder Pain Syndrome [BPS]), the bladder lining is compromised and contains lesions in advanced cases^20,21^. Based on these observations, multiple trials have been attempted to treat OAB and IC, as well as UTI, via ‘replenishment’ of the protective layer with direct instillation of complex sugar solutions into the patient’s bladder using a combination of hyaluronic acid and chondroitin sulfate ^22–24^ or hyaluronic acid alone^25^.

**Fig. 1:**
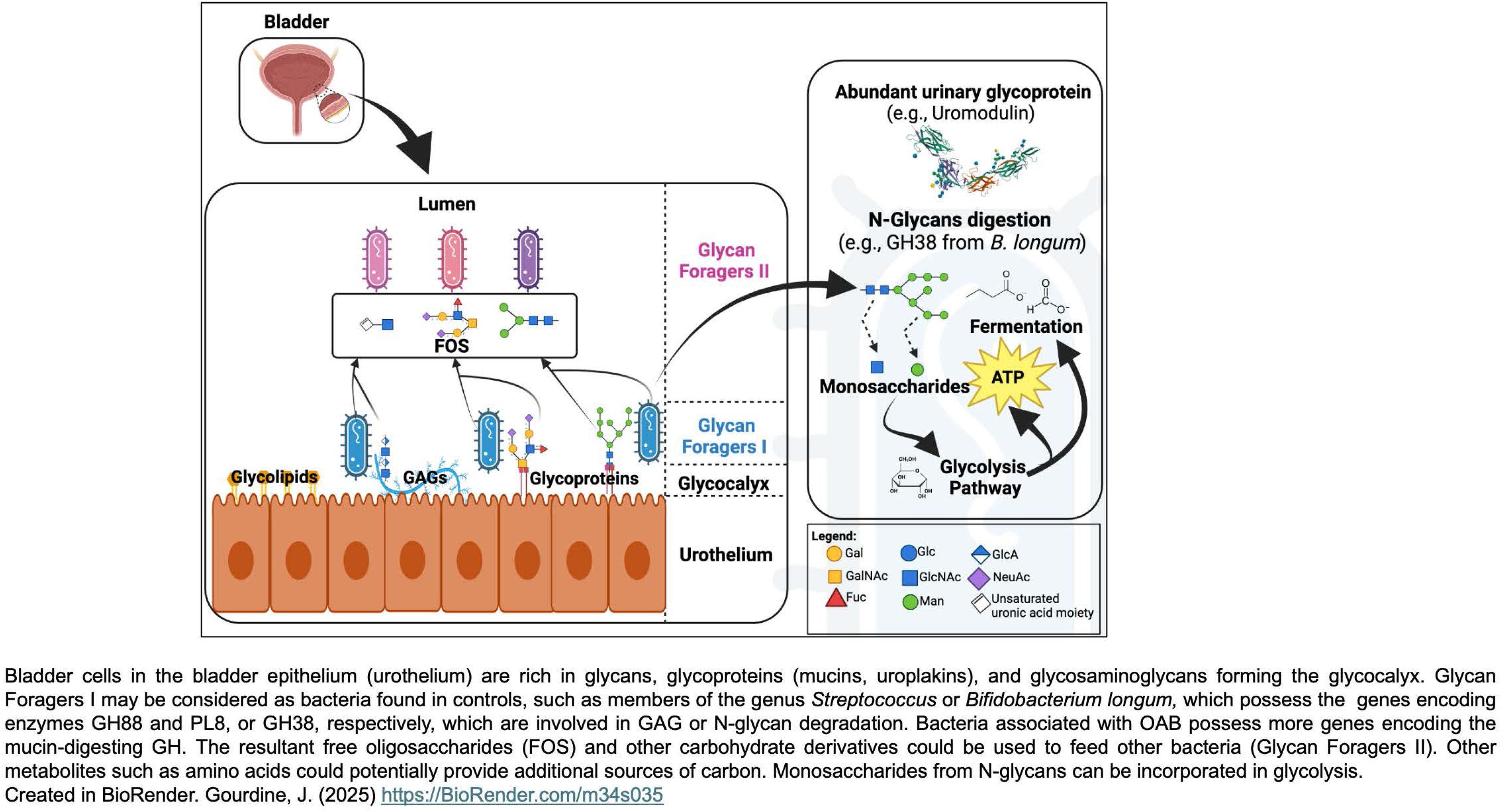
Working model of glycans utilization by the urinary microbiota.

The most abundant small metabolites in bulk human urine are urea, creatinine, hippurate, and citrate, followed by amino acids and complex sugars (glycans) and their derivatives^26^. Urea can be used for glutamine metabolism by some bacteria, such as *Proteus mirabilis*^27^, but this metabolic process can initiate apatite (stone) formation^28^. Citrate can be used as a carbon and energy source with or without co-metabolites by lactobacilli, such as *Lacticaseibacillus rhamnosus*^29^. While these two metabolites are plausible energy sources, they do not explain the whole trophic network. Indeed, many urinary bacteria cannot use these two metabolites, and a pathway that leads to stone formation is unlikely to be the main metabolic pathway for bladder bacteria. Additionally, creatinine and hippurate are unlikely to be the source of carbon for the bladder microbes since creatinine can inhibit bacterial replication^30^, and hippurate is the end product of bacterial polyphenol metabolism^31,32^. In addition to small molecules, urine contains proteins with uromodulin, constantly secreted by kidney cells, being the most abundant (∼50-100 mg/day)^33^. Uromodulin is heavily glycosylated with up to 30% of its biomass coming from N-glycans^34–37^. Indeed, uromodulin’s N-glycans are rich in mannose that permit it to be protective against uropathogenic *Escherichia coli* (UPEC), which binds to the mannose residues via its adhesin FimH^17,37^.

Combined, the bladder lining and bulk urine contain a high variety of complex sugars – glycans^26^. Unlike simple sugars or amino acids, these potential nutrients require specialized enzymes to hydrolyze them - diverse glycoside hydrolases (GHs) and polysaccharide lyases (PLs). Only microbes that produce those enzymes can benefit from these carbon sources. In turn, the presence or absence of microbial catabolism of these complex sugars can affect the functionality of the bladder protective lining, cellular communication, and urine properties. While the glycosylation pattern of bladder glycoproteins (uromodulin and uroplakins) has been shown to play a role in the adherence of bacteria and in the case of uromodulin, protection against bacteria involved in UTI^15,17,33,37^, their role in feeding commensal bladder bacteria has not been explored yet.

In other body sites rich in glycoproteins, such as the oral cavity and the gut, many members of the respective microbiome forage on host complex sugars either from glycoproteins or GAGs on host cells^38–42^, as well as free oligosaccharides (FOS) from human milk^43–46^. The bladder contains a unique repertoire of glycans and thus may select for particular microbial species and strains. There are several reports showing that microbial catabolism of sugars in the bladder is possible. Han *et al.* recently indicated the presence of unsaturated disaccharides in healthy urine, indicative of microbial GAG metabolism^47^. Additionally, De Nisco’s group showed the possible use of GAGs for selected bladder microbes from women with UTI and controls^48^, and urinary chondroitin sulfate has been found to be elevated in postmenopausal women with active rUTI^49^. In another study, Wolfe and colleagues observed differences in flocking and non-flocking strains of species in the UTI-related genus *Aerococcus* that may be related to sugar metabolism of beta-glucosides^50^.

In this study, we address the possibility that bladder bacteria digest host sugars, which may affect bladder health. We first used mass spectrometry (MS) to quantify and analyze the free soluble glycans present in urine of adult females with and without OAB to assess whether urines contain putative substrates and/or products of glycan metabolism. In both the OAB dry and UUI groups, we found similar levels of truncated glycans that may come from bacterial digestion by GHs and PLs. We then analyzed genomes of urinary bacterial isolates to identify the range of complex sugar-utilizing genes for carbohydrate active enzymes using the curated carbohydrate-active enzyme database CAZy^51^, and dbCan^52^, a Python program that that mines for GHs and PLs sequences. We then compared the presence and composition of sugar-utilizing genes in bacteria found in bladders of adult females with and without OAB.

## Material and Methods

### Urinary Free Oligosaccharides (FOS) analysis by mass spectrometry

Catheterized urine samples were collected from adult females without OAB (asymptomatic controls), with OAB but without incontinence (OAB dry), and with incontinence (OAB wet or urgency urinary incontinence [UUI]) under an IRB-approved protocol (OHSU IRB 10729) and shipped overnight on dry-ice to Emory Glycomics and Molecular Interactions Core (Emory University) for FOS analysis following the procedure described in Xia et al.^53^ Briefly, urine volume corresponding to 0.1 mg of creatinine was loaded to methanol/water-preactivated Sep-Pak C18 columns. Flowthrough was loaded onto a carbograph column preactivated with acetonitrile/ trifluoroacetate (TFA). Carbograph-bound FOS were eluted with acetonitrile/TFA (30%-0.1%), then evaporated and lyophilized. Permethylation was achieved by grinding dried FOS in NaOH in dimethylsulfoxide (DMSO) and iodomethane then dichloromethane. After centrifugation and wash steps, the lower phase was dried and resuspended in methanol. Permethylated FOS were repurified by Sep-Pak C18 columns, as described above. Eluted permethylated FOS were dried, resuspended in 50% methanol and subjected to MALDI-TOF MS using a α-dihydroxybenzoic acid (DHBA) matrix. Mass spectra were acquired by a Bruker ultrafleXtreme MALDI TOF-TOF mass spectrometer (Bruker Daltonics, Bremen, Germany) in positive ion mode with a mass range setting (m/z) of 500-5,000, and analyzed with flexAnalysis software (v3.4, Bruker Daltonics) and peaks were automatically identified with detection limits for peaks at S/N>3. (For more details, see Supplemental Methods).

### Analysis of FOS data

Post-acquisition data analysis was performed using mMass^54^. MS signals corresponding to permethylated glycans were assigned based on existing literature, public and in-house databases. For each sample, the relative abundance of each glycan compared to all glycans in the samples was calculated. Relative abundances were used for subsequent analysis and statistical purposes. Glycan annotation and assignments were based on Xia *et al.*^53^, GlyGen database^55^ (https://www.glygen.org/glycan-search), Glyconnect^56^ database, and the KEGG database^57^, and represented following the Symbol Nomenclature for Glycans (SNFG)^58^ using the Glycoglyph^59^ interface (https://glycotoolkit.com/Tools/GlycoGlyph/). FOS were organized according to their features (terminal galactose, fucosylated, mannosylated, sialylated, glucose polymers, or a combination) for further analysis.

### Genes encoding Glycoside Hydrolases (GH) and Polysaccharide Lyases (PL) from urinary bacterial genomes

Bacterial genomes isolated from adult females with and without OAB were selected from publicly available data originating from Thomas-White et al.^60^. Sequence Read Archive (SRA) and the European Nucleotide Archive (ENA)^61,62^ under the accession number PRJEB8104; See Supplemental Table 2). GH and PLs were identified from the selected urinary metagenomes using the Carbohydrate Active Enzyme Database (CAZy)^63,64^ and the meta server dbCAN2^52^ on Python^65^ and Linux with the default parameters and quantified in R^66^ version 4.2. dbCAN2 uses three CAZyme annotation tools: Hotpep^67^, HMMER^68^, and DIAMOND^69^. If 2 or more databases returned the same gene for a genome, it was counted as an identified GH or PL gene. Potential substrates for the GH/PL family were attributed manually based on curated spreadsheet from dbCAN2^52^ (last updated 08/25/2022) CAZy database^63,64^, Berlemont and Martini ^70^, Kaoutari *et al.*^71^ and Tailford *et al.*^39^. Two groups of potential substrates were defined based on a broad-level classification of glycan molecules (e.g., host glycans) and on a more granular-level classification of glycan molecules (e.g., N-glycans, mucin, GAGs).

### Statistical analysis

Statistical analyses were performed in R. Differences between groups in urinary free oligosaccharides abundances were assessed using Kruskal-Wallis tests. For each glycan, groups were compared using a one-way ANOVA (analysis of variance) test. A post-hoc analysis (t-test) was performed using a correction for multiple comparisons. Statistical differences in the presence of GH and PLs between control and OAB were evaluated with the Fisher’s exact test.

## Results

### Analysis of urinary free oligosaccharides in OAB and control cases

Thirteen out of sixty samples yielded a low signal to noise ratio or low intensity peaks and were not used for further analysis. The average FOS counts for the remaining 47 samples was 82 (+/- 24.9) with mass spectra ranging from 500.5to 3780.3 m/z. Fifty-eight urinary FOS of superior quality (relative intensity >0.1) across the samples were further annotated using Glycoglyph^59^, Glygen database^55^ and curated literature^53^. An exemplary of an annotated urinary FOS profile by mass spectrometry (S28) is given in Fig. 2A.

**Fig. 2:**
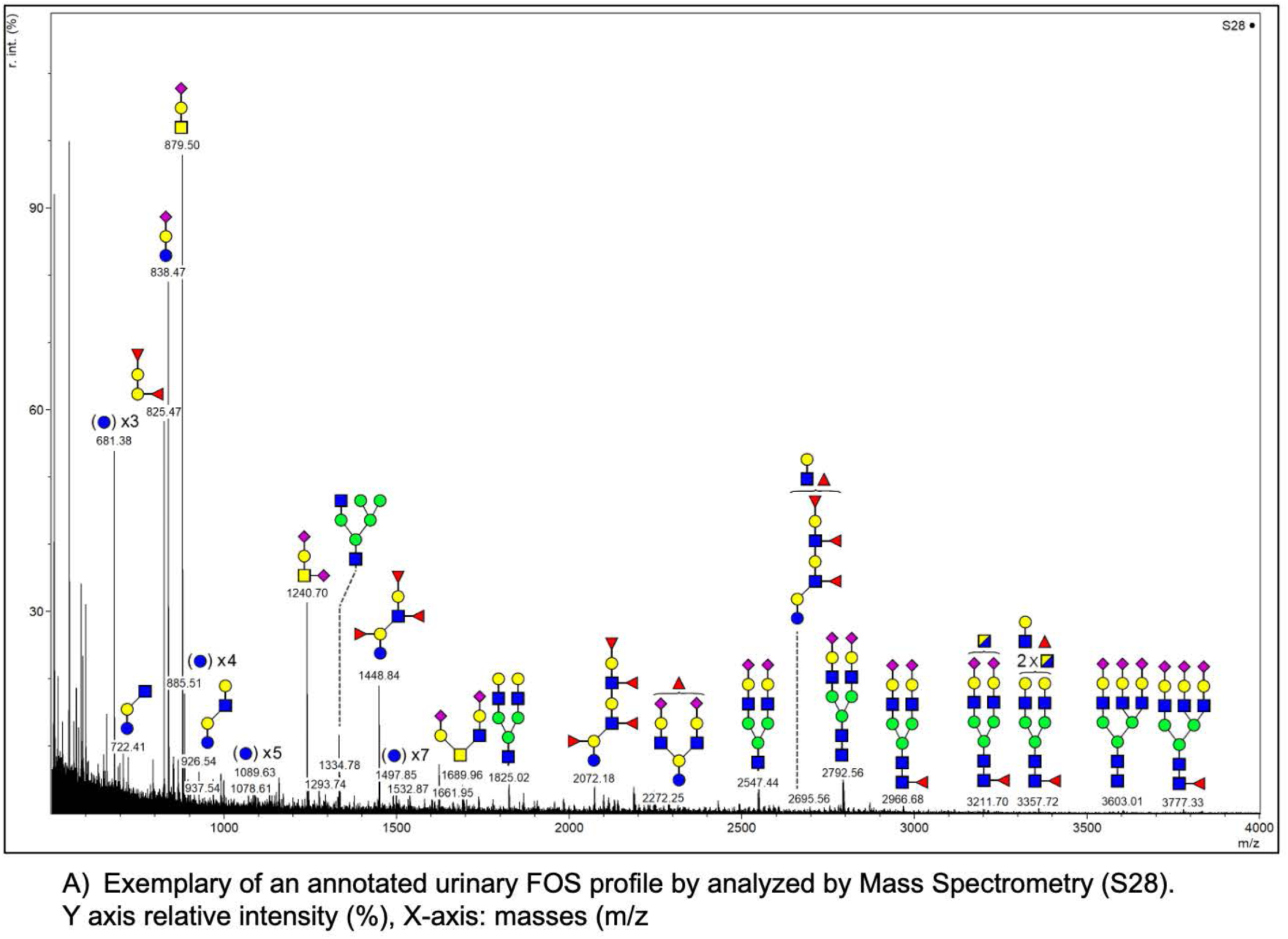
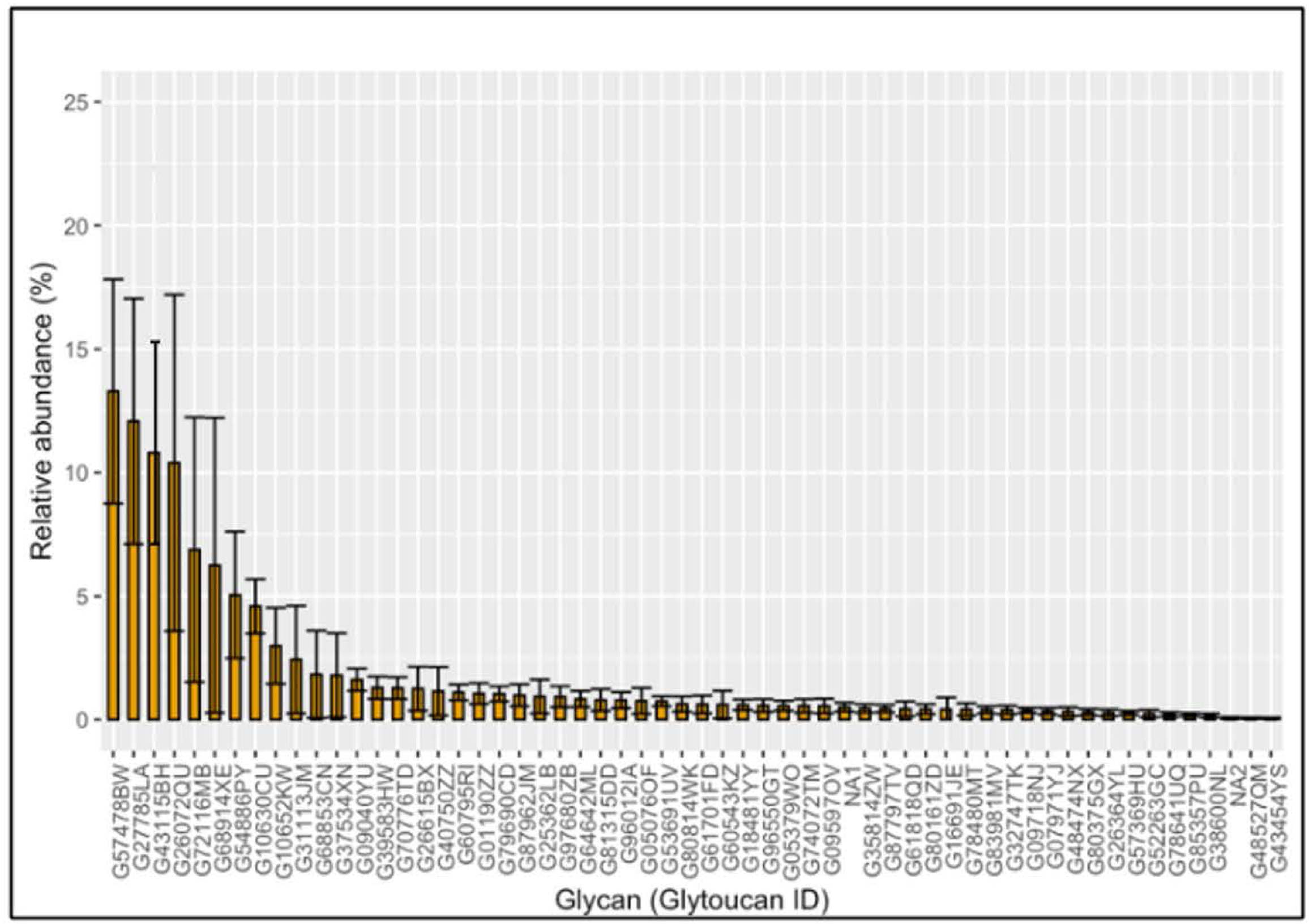
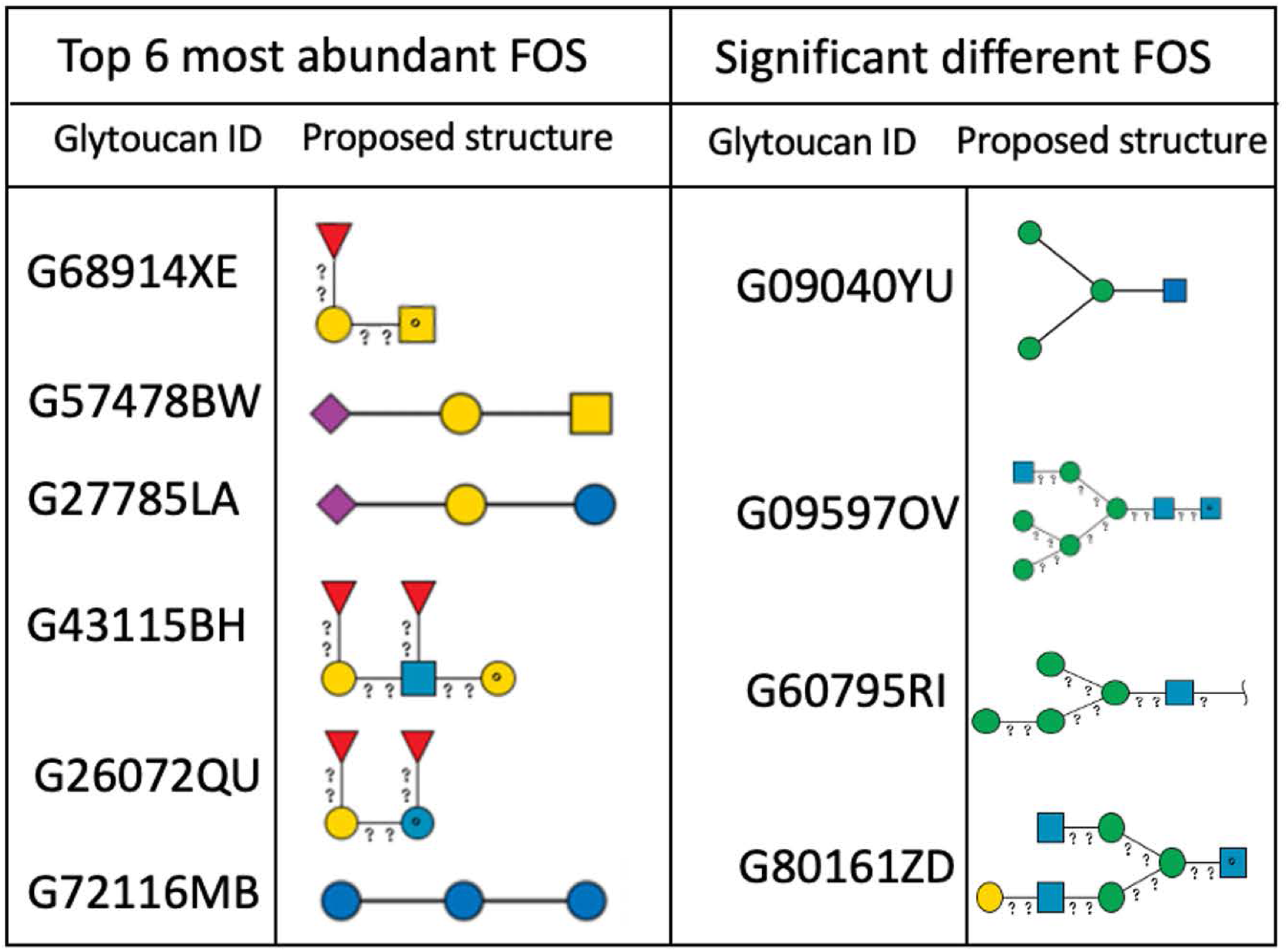
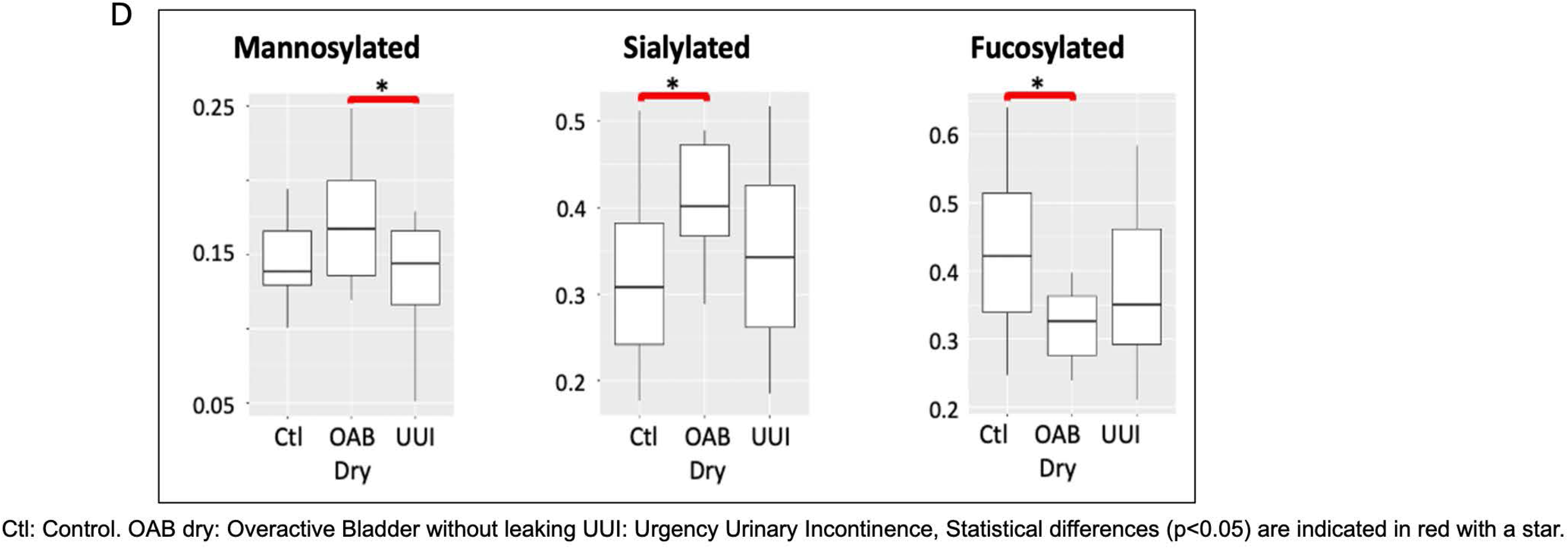
Urinary FOS. A) Exemplary of an annotated urinary FOS profile by analyzed by Mass Spectrometry, B) Distribution of FOS in urine from women with UUI, C) Representative structures of FOS in urine from women with UUI, D) Statistically different FOS between control, OAB Dry, and UUI, grouped by terminal feature

We identified GlyTouCan IDs for 56 of the 58 selected FOS (Fig. 2B and Supplemental Table S1). The two exceptions were NA1 (m/z= 1601.9 and composition: Hex1HexNAc4Fuc2) and NA2 (m/z= 3357.9, composition: Hex6HexNAc7Fuc2). Overall, urinary FOS were characterized by a low relative abundance (<25%). Regardless of participant group and ranking, the seven most abundant FOS (average relative abundance ∼5-15%) have low molecular weights (m/z between 500-900) and were identified as fucosylated Gal-GalNAc (GlyTouCan ID: G68914XE), sialyl Gal-GalNAc (GlyTouCan ID: G57478BW), sialyl-lactose (GlyTouCan ID: G27785LA), difucosylated lactosamine (GlyTouCan ID: G43115BH), difucosylated lactose (GlyTouCan ID: G26072QU) and a triglucose (GlyTouCan ID: G72116MB) (Fig. 2C).

To determine if glycosylation pathways differed between cohorts, we regrouped FOS by glycosylation features (e.g., mannosylated, fucosylated, sialylated, and polyglucose glycans). Statistically significant differences were observed for mannosylated, sialylated, and fucosylated glycans, respectively, between patient groups (Table 1, Fig. 2D). Between control and OAB dry, four individual FOS differed significantly; two mannosylated, one sialylated, and one fucosylated. Among them, two FOS were higher in OAB dry (one mannosylated and one sialylated, GlyTouCan ID: G09040YU, G277785LA). Two FOS were lower (one mannosylated and one fucosylated, GlyTouCan ID: G6079RI, G70776TD). Between UUI and OAB dry, six FOS differed significantly, three mannosylated and three fucosylated. Three mannosylated FOS were higher in OAB dry (GlyTouCan ID: G09040YU, G80161ZD, G09597OV) and three fucosylated FOS were lower in OAB dry (GlyTouCan ID: G70776TD, G48474NX, G40750ZZ). No differences were identified between controls and UUI.

**Table 1:**
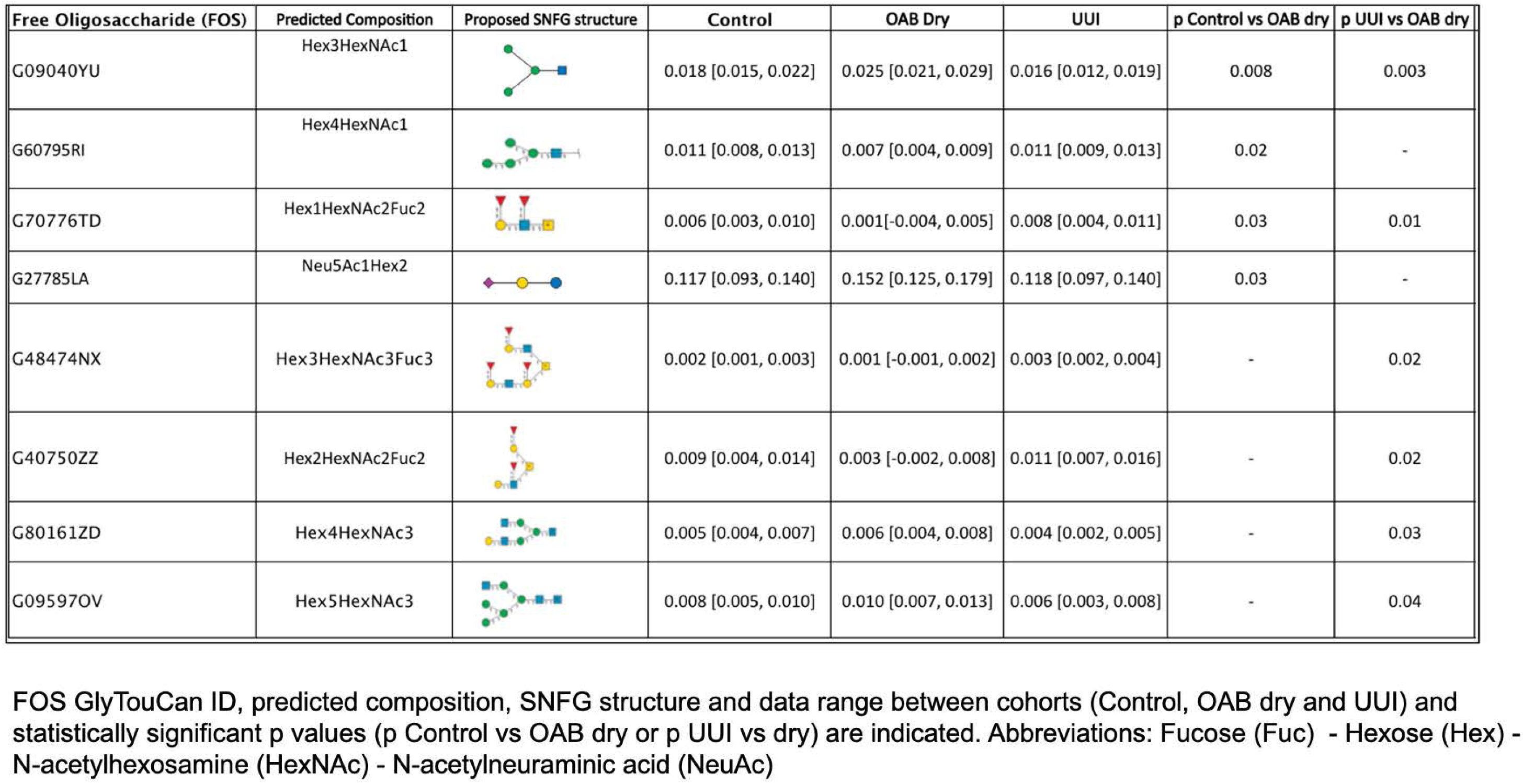
Statistically significant different urinary FOS between OAB dry, UUI and control.

| Free Oligosaccharide (FOS) | Predicted Composition | Proposed SNFG structure | Control | OAB Dry | UII | p Control vs OAB dry | p UII vs OAB dry |
| --- | --- | --- | --- | --- | --- | --- | --- |
| G09040YU | Hex3HexNAc1 |  | 0.018 [0.015, 0.022] | 0.025 [0.021, 0.029] | 0.016 [0.012, 0.019] | 0.008 | 0.003 |
| G60795RI | Hex4HexNAc1 |  | 0.011 [0.008, 0.013] | 0.007 [0.004, 0.009] | 0.011 [0.009, 0.013] | 0.02 | - |
| G70776TD | Hex1HexNAc2Fuc2 |  | 0.006 [0.003, 0.010] | 0.001[-0.004, 0.005] | 0.008 [0.004, 0.011] | 0.03 | 0.01 |
| G27785LA | Neu5Ac1Hex2 |  | 0.117 [0.093, 0.140] | 0.152 [0.125, 0.179] | 0.118 [0.097, 0.140] | 0.03 | - |
| G48474NX | Hex3HexNAc3Fuc3 |  | 0.002 [0.001, 0.003] | 0.001 [-0.001, 0.002] | 0.003 [0.002, 0.004] | - | 0.02 |
| G40750ZZ | Hex2HexNAc2Fuc2 |  | 0.009 [0.004, 0.014] | 0.003 [-0.002, 0.008] | 0.011 [0.007, 0.016] | - | 0.02 |
| G80161ZD | Hex4HexNAc3 |  | 0.005 [0.004, 0.007] | 0.006 [0.004, 0.008] | 0.004 [0.002, 0.005] | - | 0.03 |
| G09597OV | Hex5HexNAc3 |  | 0.008 [0.005, 0.010] | 0.010 [0.007, 0.013] | 0.006 [0.003, 0.008] | - | 0.04 |

FOS GlyTouCan ID, predicted composition, SNFG structure and data range between cohorts (Control, OAB dry and UII) and statistically significant p values (p Control vs OAB dry or p UII vs dry) are indicated. Abbreviations: Fucose (Fuc) - Hexose (Hex) - N-acetylhexosamine (HexNAc) - N-acetylneuraminic acid (NeuAc)

### GH/PL abundance in bladder microbiomes of OAB and control groups

We analyzed genomes from 111 bacterial strains from the Thomas-White et al. dataset^60^. From adult females with OAB (not further classified into OAB dry and UUI), we analyzed 67 strains from 47 species; from asymptomatic controls, we analyzed 44 strains from 31 species. As indicated in Fig. 3A, the genes for 80 GHs/PLs were predicted in bladder bacteria from the OAB and control groups, with GHs predominant (∼90%). The most common GHs predicted from both datasets (>50%) were GH13, GH77, GH3, GH32, GH1, and GH2, while the least abundant, common in both control and OAB cohorts, were GH39, GH79, and PL22. Some GH/PL families were unique to the OAB dataset (GH108, GH17, GH50, GH84, GH120, GH19 and PL4, PL5, PL7) or to the control dataset (GH121, GH129, GH144, GH26, GH27 and PL17).

**Fig. 3:**
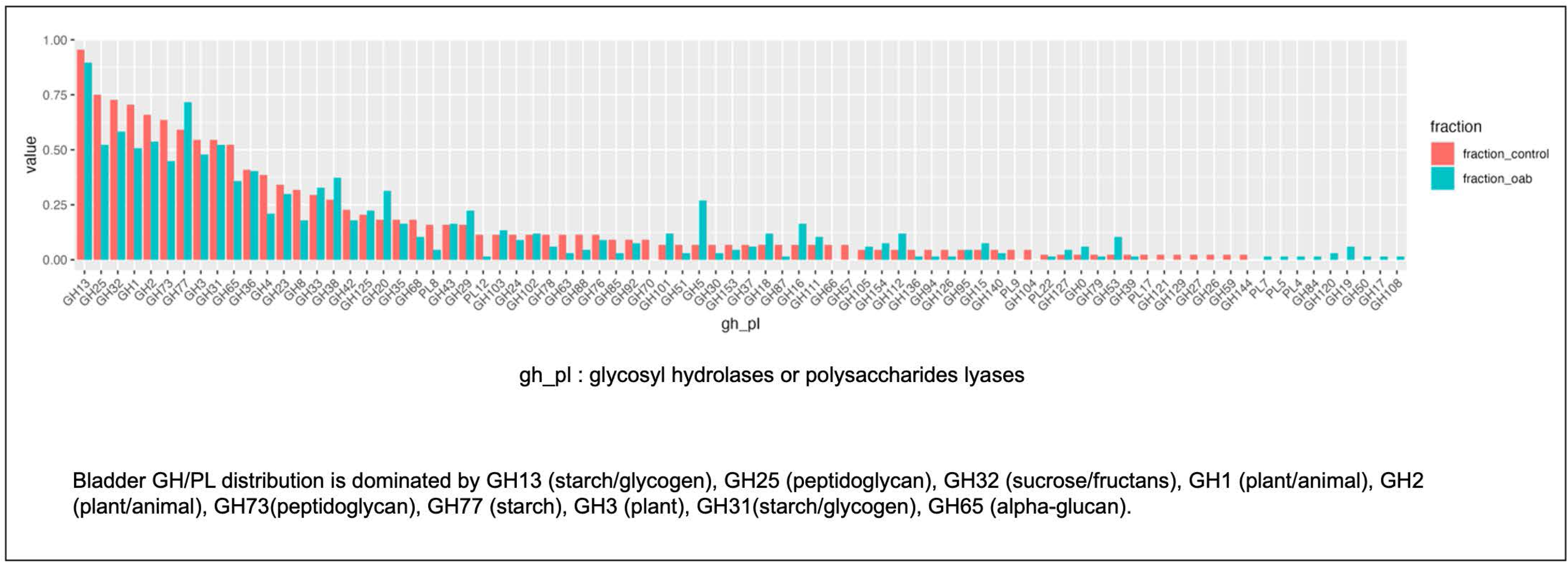
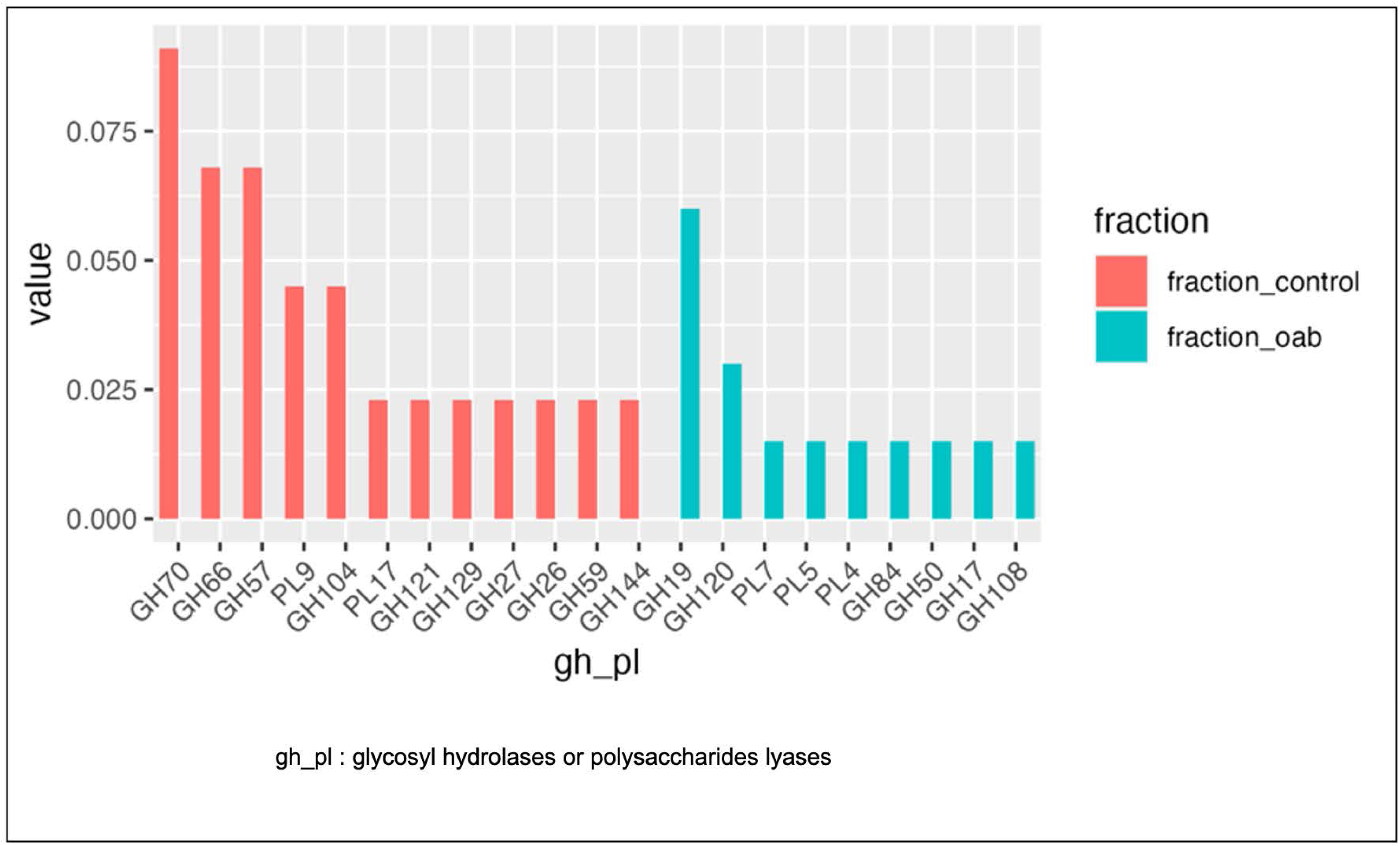
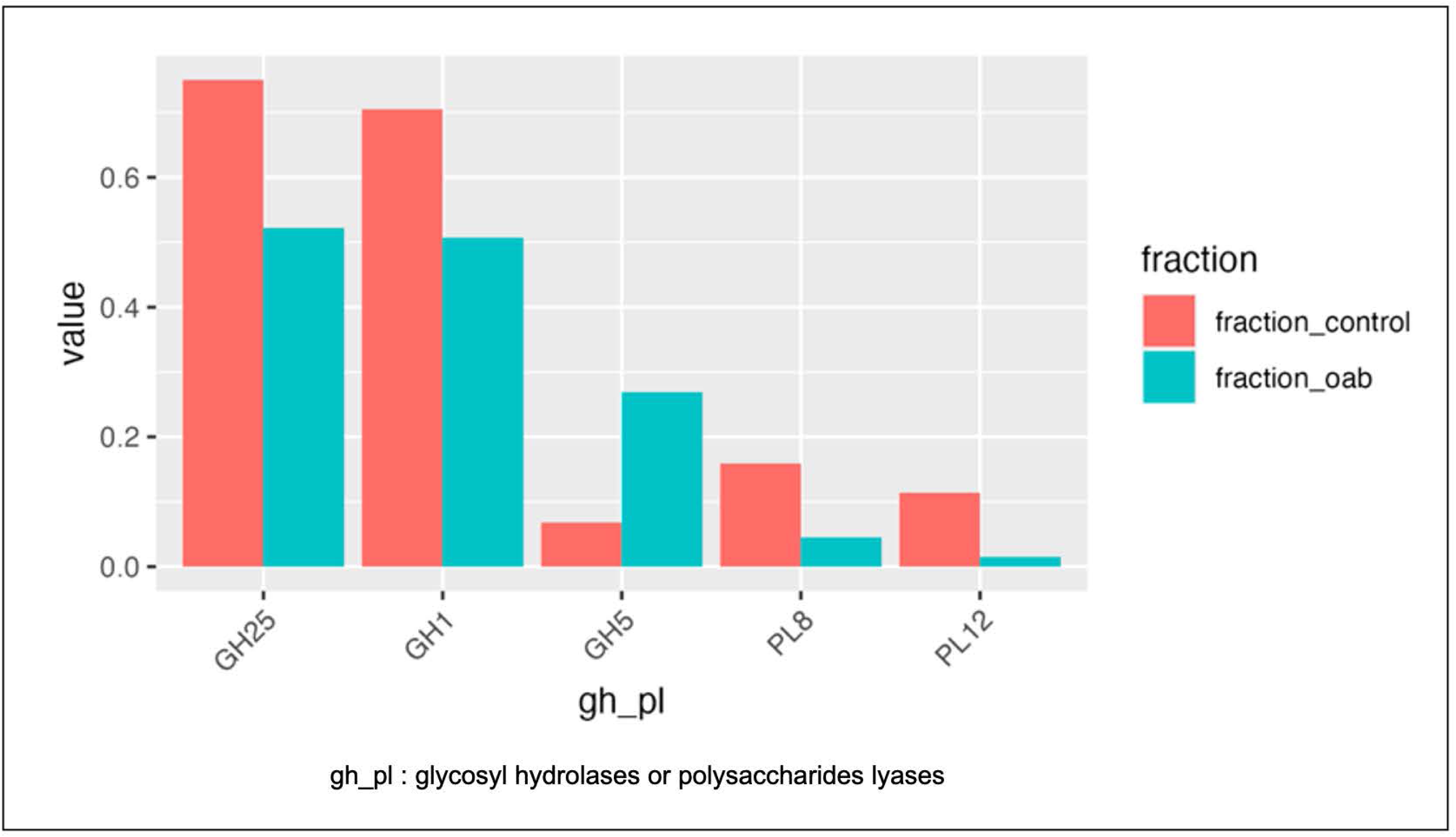
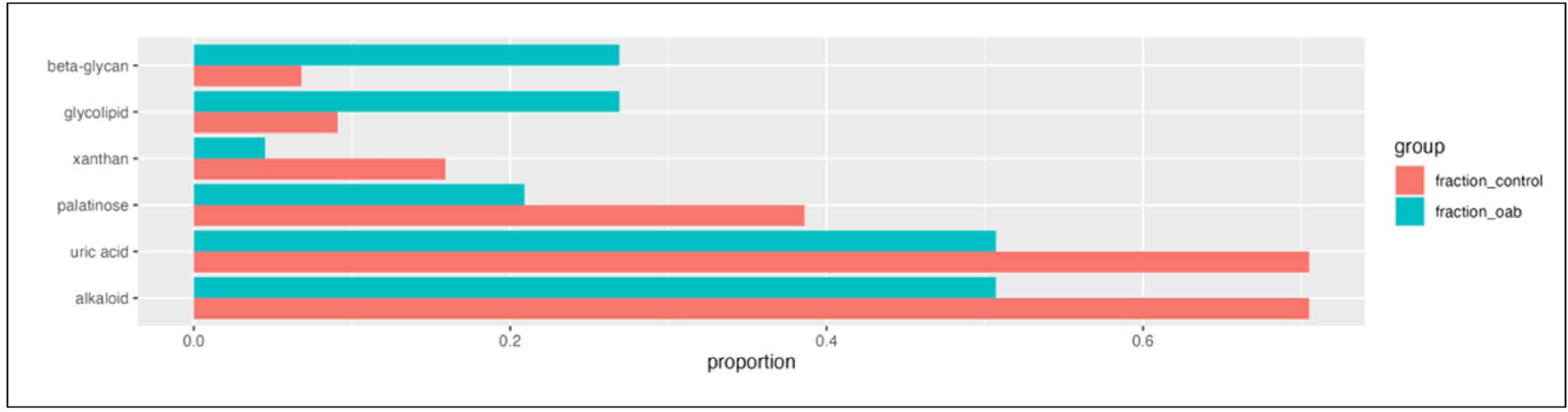
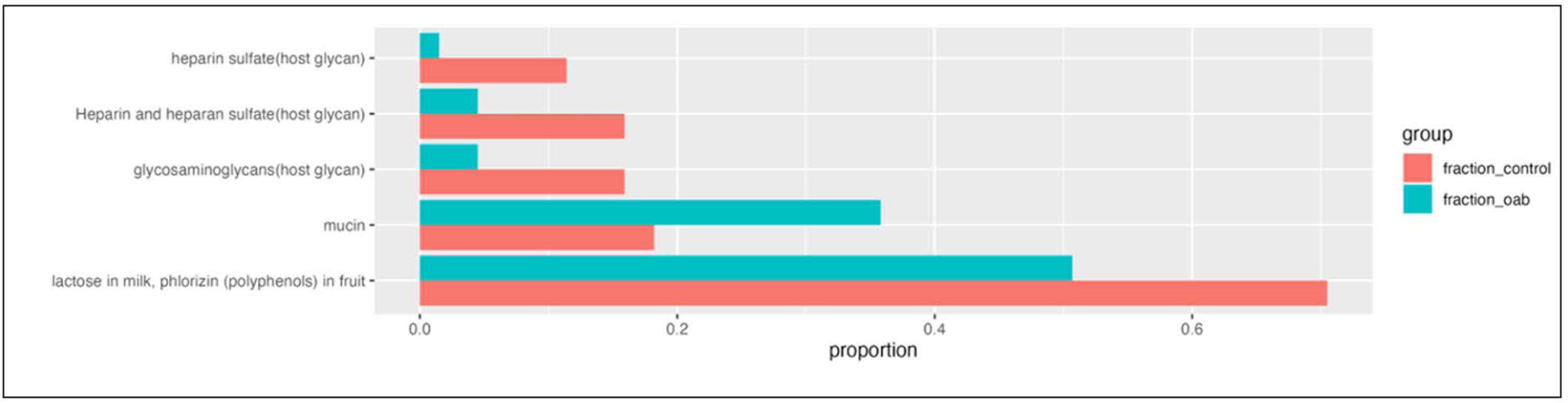
Glycoside hydrolase and polysaccharides lyases from bladder bacterial isolates. A) Bladder GH/PL distributions, B) Unique GH / PL in control and OAB bladder bacteria, C) Statistically significant GH/PL in OAB and control, D) Statistically significant potential broad-level substrates for GHs/PLs in OAB and Control, E) Statistically significant potential granular-level substrates for Bladder Microbiome enzymatic capacity in OAB and Control

### Bladder microbiomes of both, OAB and control, groups have genes encoding unique GHs and PLs

As indicated in Fig. 3B, the genes for 12 GHs/PLs were more unique in the control group, with potential substrates as follows: alpha-glucan (GH70, GH66, GH57), starch (GH57), peptidoglycan (GH104), pectin (PL9, GH129), alginate (PL17), arabinan (GH121), host glycan mucin-o-type O-glycans (GH129, GH27), raffinose (GH27), alpha-galactan (GH27), beta-mannan (GH26), xylan (GH26), beta-glucan (GH26, GH144), beta-mannan (GH59), glycolipids (GH59). For OAB, 9 GHs/PLs were unique with the following potential substrates chitin (GH19), peptidoglycan (GH19, GH108), arabinan (GH120), host glycan – alginate (PL5), alginate (PL7), host glycan - hyaluronan (PL7, GH84), host glycan-mucin (GH84), pectin (PL4), alginate (PL4), arabinogalactan (PL4), agarose (GH50), beta-glucan (GH17).

### Statistically significant GHs/PLs in bladder microbiome of OAB and control groups

The genes for 5 GHs/PLs were statistically significant (p-value <0.05) between the OAB and control groups (Fig. 3C). GH25, GH1, PL8 and PL12 were more abundant in controls, while GH5 was more abundant in the OAB group. Host GAGs (PL8 and PL12), peptidoglycans (GH25), and beta-glucan (GH5, GH1) are potential substrates for these enzymes (Fig. 3D).

### Potential substrate of GHs/PLs in bladder microbiome in OAB and control groups

To examine common and disparate glycan utilization between the bladder microbiota of women with and without OAB, we compared the potential substrates for the predicted GHs and PLs (Fig. 3D, Supplementary Fig. S1). We identified several substrates with statistically significant (p-value <0.05) differences between groups based on the classifications of glycan molecules either at the broader level (e.g., host glycans, Fig. D) and on a more granular level (e.g., mucins from host glycans, Fig. 3E). Substrate glycans associated with alkaloid, uric acid, palatinose and xanthan were higher in controls, while beta-glycan and glycolipids were higher for the OAB group. Even though uric acid is not a glycan, its degradation involves hydroxyisourate hydrolase, which possesses sequence homology with many GH^72^. Many alkaloids include glycan moieties and their metabolism uses GH^73^. To explore potential differences between host glycans utilization, we compared host substrates based on GHs/PLs specificity (Supplementary Fig. S2). Bacterial strains from controls indicated more putative enzyme using substrate heparan (GAG), while genes encoding enzymes digesting mucin-type glycans were more represented in bacterial strains from the OAB group (Fig. 3E).

### Glycoside hydrolases and polysaccharides lyases in genera found in both OAB and control

We identified several differences in the GHs and PLs present in 12 genera with isolates originating from both control and OAB cohort patients (Fig. 4). Unique GH/PL features are found for the genera *Streptococcus* and *Schaalia* for controls and *Actinomyces* for OAB. For *Streptococcus*, unique GHs/PLs were the following with potential substrates indicated in parenthesis: PL8 (host glycans, GAGs), PL12 (host glycans, GAGs), GH101 (host glycan, mucin), GH78 (plant polysaccharides), GH88 (host glycans, GAGs), GH87 (fungal polysaccharides), GH68 (plant polysaccharides), GH79 (host glycans, GAGs), GH70 (plant polysaccharides), and GH66 (alpha-glucan). Some of the common features for these unique streptococcal GHs/PLs are host glycans utilization, particularly for GAGs (e.g., PL8, PL12, GH88, GH79). For *Actinomyces*, unique GHs/PLs are the following with potential substrates indicated in parenthesis: GH73 (peptidoglycans), GH4 (oligosaccharides), GH101 (host mucin), GH127 (plant polysaccharides), GH51 (plant polysaccharides), GH20 (host glycan mucin), GH125 (plant N-glycan), GH5 (plant polysaccharides), GH38 (host glycan, fungal polysaccharides), GH30 (host glycan, plant polysaccharides), GH78 (plant polysaccharides), GH63 (host glycan, plant polysaccharides), GH18 (host N-glycan, peptidoglycans), and GH120 (plant polysaccharides).

**Fig. 4:**
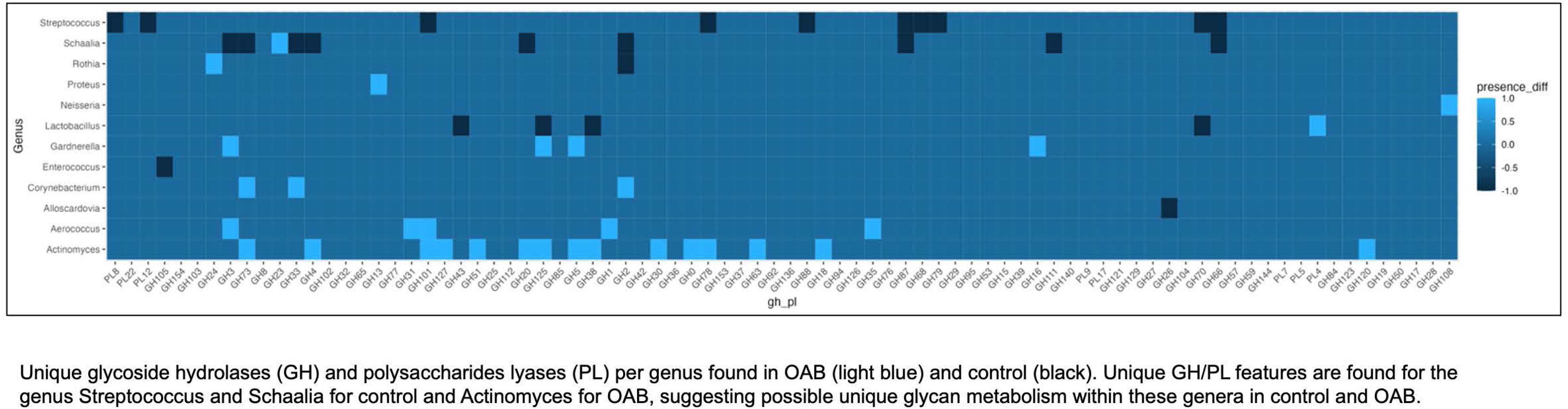
Heatmap of unique glycoside hydrolases and polysaccharides lyases according to common genus found in OAB and control.

## Discussion

### FOS diversity and differences as potential markers of OAB

Urinary FOS have been used as disease markers for host glycan metabolism defects, as well as many types of cancers^53,74^ but have not been investigated in the context of OAB. In this study, urinary FOS were characterized by a low relative abundance (<25%) with sialyl-lactose, sialylated Gal-GalNAc, fucosylated lactose and derivative, and triglucose being the most abundant FOS (Fig. 2C). These results are comparable to published literature for healthy individuals’ urine (Xia et al. ^53^, and Hanzawa et al.^75,76^).

Our data indicated that four FOS are significantly different in women with dry OAB differentiating this cohort from both other cohorts - control and UUI (Fig. 2CD, Table 1). These four FOS may come from either N-glycans, as they all share a core tri-mannose fragment, or originate from N-glycosylated uromodulin that often contains similar fragments. The most abundant of these four differentially present FOS is tri-mannosyl-GlcNAc that thus could be used as a putative marker of OAB dry condition.

### Evidence for microbial catabolism of polysaccharides in the urinary tract

In both the OAB and control groups, the catheterized urine samples yielded a high diversity of glycans. Based on the predicted structures, these FOS may come from glycoproteins (N and O-glycans), glycolipids, and/or glycogen (Table 2). Based on the bladder ultrastructure, we can hypothesize that some polysaccharides come from the N-linked glycoproteins (uroplakins and uromodulin) and the O-linked glycoproteins (mucins). Other polysaccharides may come from degradation and recycling of other diverse glycoproteins and glycolipids during the shedding of urothelium cells and some low molecular weight urinary FOS come from diet.

**Table 2:**
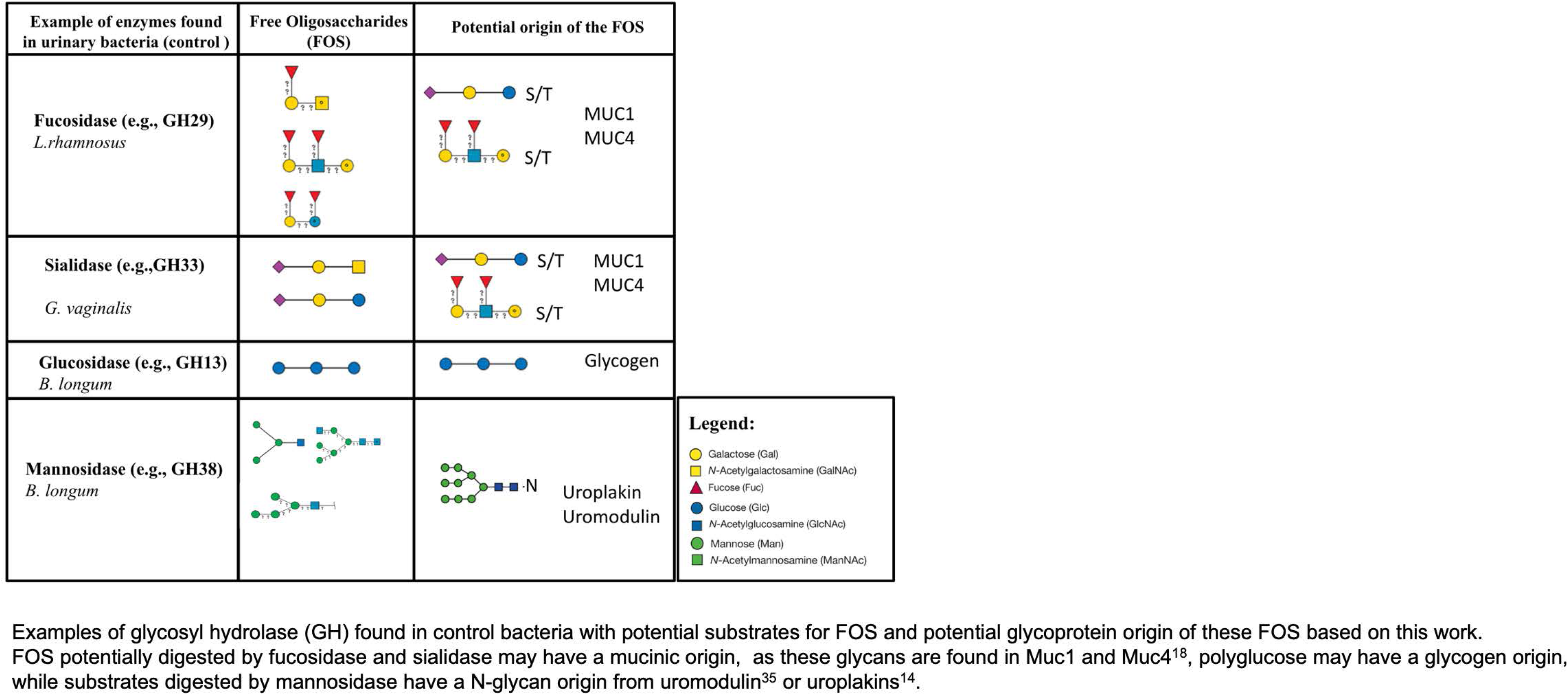
Examples of glycosyl hydrolase (GH) found in control bacteria with potential substrates for FOS.

| Example of enzymes found in urinary bacteria (control ) | Free Oligosaccharides (FOS) | Potential origin of the FOS |
| --- | --- | --- |
| **Fucosidase (e.g., GH29)**<br>*L.rhamnosus* |  | S/T MUC1<br>S/T MUC4 |
| **Sialidase (e.g.,GH33)**<br>*G. vaginalis* |  | S/T MUC1<br>S/T MUC4 |
| **Glucosidase (e.g., GH13)**<br>*B. longum* |  | Glycogen |
| **Mannosidase (e.g., GH38)**<br>*B. longum* |  | Uroplakin<br>Uromodulin |

- Galactose (Gal) - N-Acetylgalactosamine (GalNAc) - Fucose (Fuc) - Glucose (Glc) - N-Acetylglucosamine (GlcNAc) - Mannose (Man) - N-Acetylmannosamine (ManNAc)
Examples of glycosyl hydrolase (GH) found in control bacteria with potential substrates for FOS and potential glycoprotein origin of these FOS based on this work. FOS potentially digested by fucosidase and sialidase may have a mucinic origin, as these glycans are found in Muc1 and Muc4<sup>18</sup>, polyglucose may have a glycogen origin, while substrates digested by mannosidase have a N-glycan origin from uromodulin<sup>35</sup> or uroplakins<sup>14</sup>.

Bladder cell surface and secreted glycoproteins (uroplakins and uromodulins) have high mannose N-glycans^15,36^. However, our FOS data showed a low abundance of free high mannose glycans. This might indicate that host N-glycans are microbially digested, hence resulting in the observed absence of high mannose FOS and the presence of truncated mannosylated FOS. This serves as indirect evidence that at least some urinary FOS come from bacterial degradation of urinary glycoproteins. In fact, *Enterococcus faecalis* and *Bifidobacterium longum* species are found in the urobiome^60^ and strains of these species isolated from other microbiomes have been reported to digest high-mannose containing glycoproteins (e.g., RNase B) *in vitro*^77,78^.

Similarly, the presence of polyglucose of different molecular weights (from triose to pentaglucose), in combination with the presence of the GH13 gene in many bacterial genomes from both OAB and controls, suggests utilization of host glycogen by the urobiome, similar to the digestion of glycogen in the vaginal tract by *Lactobacillus vaginalis, Lactobacillus crispatus and Gardnerella spp* ^79–81^. Urinary polyglucose has been reported in healthy conditions^53,82^. Their origins may be diverse, for instance urinary tetraglucose may come from a carbohydrate-rich diet or liver/muscle glycogen metabolism^83^.

Finally, presence of the trisaccharides sialyl-lactose and sialyl-Gal-GalNAc can be correlated to microbial GH33 enzyme genes from both the OAB and control groups, possibly indicating digestion of sialic acid O-glycans by the urobiome. This would be similar to vaginal isolates of *Prevotella* species and *Gardnerella vaginalis* that were observed to digest sialic acid from vaginal mucins^84^. Interestingly, in the first urinary metagenome-genome-wide association of Chinese individuals, Zou et al. reported a host genetic variant of B4GALNT2 (rs202106085) that may influence the urobiome^85^; B4GALNT2 encodes a glycosyltransferase involved in the addition of GalNAc to sialyl-Gal (Blood group Sd/Cad antigen) to O-glycoprotein like mucin^86^. Hence, mucin glycosylation may be relevant to explore in the urobiome context.

### Changes in GHs/PLs composition in species and strains in OAB cohort

Similar to the microbiota associated with other body sites (e.g., gut, oral cavity), glycoside hydrolases of bladder microbiota are predominated by families involved by energy production such as GH13, GH32, GH1 and peptidoglycan breakdown (GH25, GH73)^64,87^. Contrary to the vaginal microbiota, in which GH4, GH68, GH42 and GH8 are more prevalent, predominant GHs/PLs of the urinary microbiota are characterized by their potential substrates originating from diet (e.g., glucan), as well as glycogen.

While some GHs and PLs are unique in datasets from the OAB and control groups, they seem to converge toward the same broad substrate types for both the OAB group (i.e., mucin [GH84] and plant/algal [GH50, PL4, PL5, PL7]) and for control (i.e., mucin [GH129], plant/algal [GH121, GH144, GH26, GH17], and oligosaccharides [GH27])^64,87^. These data suggest that in both groups, bladder microbes may have some capacity to digest host mucins as well as dietary plant/algal carbohydrates (e.g., fructan).

Three families of GHs were unique to OAB with the following potential substrates: peptidoglycan (GH108), broad glucan (GH17) and chitin (GH19). Control fractions have more genes predicted to encode enzymes that can potentially digest GAGs, particularly heparan, compared to OAB cases. For instance, PL8 in *Bacillus* sp. UMB0893, *L. rhamnosus*, and *Streptococcus agalactiae* found uniquely in our control set. On contrary, PL8 was present in *P. mirabilis* and *E. faecalis* from both control and OAB groups. Urinary isolates of *P. mirabilis* and *E. faecalis* have been shown to degrade GAGs *in vitro* in pure cultures^48^, confirming this part of our analyses. As indicated in Fig. 4, members of the genus *Streptococcus* are predicted to possess unique GHs/PLs with host GAG utilization while members of the genus *Actinomyces* are predicted to possess more GHs/PLs involved in host mucin and plant carbohydrates degradation. OAB fraction possesses more bacteria with GH that may be involved in host mucin and glycolipids digestion (Fig. 3E).

Our data also show some GHs/PLs differences at the strain level. For instance, GH5 genes were found in 5 strains of *G. vaginalis* from OAB cohort while absent in two other OAB strains and absent in three strains isolated from control group. These granular differences at the strain level could explain the reason why it has been hard to differentiate bacterial OAB markers using bacterial composition, which is typically limited to genus or species level analyses.

### Study limitations

Our study has several limitations that warrant more in-depth follow-up studies. Due to the MS methodology, we could not detect simpler sugars, such as unsaturated disaccharides. In our bioinformatic analysis, we focused on GH/PL genes and thus might have missed catabolic enzymes with broader spectrum of glycan-binding domains, such as other esterases. Next steps will include more functional studies to test our model, including glycan metabolic labelling and metatranscriptomics, as has been done for other bodies sites^88^

Our genomic analyses show the presence of GH enzymes usually involved in plant and algal carbohydrate digestion. A careful look at patients’ diet may also help identify potential glycans coming from food (e.g., inulin), as certain dietary oligosaccharides and monosaccharides (e.g., mannose), can end up in the bloodstream, in the GI, and in the urinary tract^89^. In this study, we did not control for the diet of patients but, in the future, it will be of interest to investigate how dietary complex carbohydrate can affect the urobiome composition. Lastly, our analysis has some limitations due to the sample size, and future work includes expanding the analysis to larger cohorts.

### Conclusions

Our study is the first attempt to connect free urinary glycans to bladder microbial metabolism in OAB. Four urinary glycans were different between OAB dry and control. Overall, our results show significant difference in 9 GH/PL between bacteria coming from bladder of OAB versus healthy women. Analyzing putative substrates of the identified GH/PL enzymes we predicted overlapping GH/PL activities in the bladder microbiota regardless of differences between GH/PL in bacteria originating from women with OAB and controls and showing that in general, bladder bacteria can digest mucins O-glycans, N-glycans and free oligosaccharides in the urine. We also identified that bladder microbiota has the potential ability to digest host GAGs, N-glycans and O-glycans. Statistically significant differences between OAB and control groups in potential for the host glycan utilization were identified, suggesting a higher utilization of GAGs in control cohort compared to OAB cohort.

## Acknowledgments

The authors acknowledge the Circle of Giving from the Center for Women’s Health at OHSU (LK and JPG), the M.J. Murdock Charitable Trust at Lewis & Clark College (JPG), and the National Institutes of Health (NIH) for support of this project through awards NIDDK K01DK116706 (LK), NCATS UL1TR002369 (LK), NCATS UL1TR002369-03S1 (JPG).

The authors also thank the Oregon Clinical Translational Institute (OCTRI) especially Dr. David Ellison and Chris Larsen for their support, Amanda Holland (OHSU) and Sean Davin (OHSU) for assistance with sample and data management, Dr. Sylvain Lehoux for assistance with glycan analysis, and Yi Lacenkanek and Dr. Xuesong Xue at Emory University.

This study was also supported in part by the Emory Glycomics and Molecular Interactions Core (EGMIC), which is subsidized by the Emory University School of Medicine and is one of the Emory Integrated Core Facilities. Additional support was provided by the National Center for Advancing Translational Sciences of the National Institutes of Health under Award Number UL1TR002378.

The content is solely the responsibility of the authors and does not necessarily reflect the official views of the National Institutes of Health or any other funder.

## Abbreviations

OAB: Overactive Bladder
UUI: Urgency Urinary incontinence
FOS: Free oligosaccharides
GH: Glycoside Hydrolase
PL: Polysaccharide Lyase
GlcNAc: N-acetylglucosamine
GalNAc: N-acetylgalactosamine
Gal: Galactose
Fuc: Fucose
Hex: Hexose
HexNAc: N-acetylhexosamine

---

**Supplemental Table1. Urinary FOS in Bladder Microbiome annotation and potential GHs/PLs utilization**

Tab 1: Samples label

Tab 2: Data sorted by group (m/z, relative intensity)

Tab 3: Final selection of samples (m/z, relative intensity)

Tab 4: Composition of the glycans

m/z, composition (Hex, HexNAc, Fuc, NeuAc), Predictive Composition, Proposed GlyTouCan ID, Potential GH/PL, Potential substrate, references, percent of relative intensity and Standard Deviation (Std_Dev)

Tab 5: Heat Map of the FOS according to OAB Tab 6: Analysis by composition groups

Tab 7: ANOVA per group based on the type of glycans (Fucosylated, sialylated, glucose polymers

Tab 8: ANOVA per glycan where significant glycans are highlighted in green

Tab 9: Post hoc with Bonferroni correction for significant FOS

**Supplemental Table 2. Bacterial isolates used from the analysis from Thomas-White et al.**

Tab 1: Bacterial_Isolates, Bacterial isolates used from the analysis from Thomas-White et al. Assembly, Level, WGS, Biosample, Strain, Taxonomy and Group (OAB or Control) were indicated.

Tab 2: Control_GH_PL, Bacterial isolates and GH/PL counts for controls Tab 3: OAB_GH_PL, Bacterial isolates and GH/PL counts for OAB

**Supplemental data: Raw files Mass spec files**

**Supplemental Fig.1.**
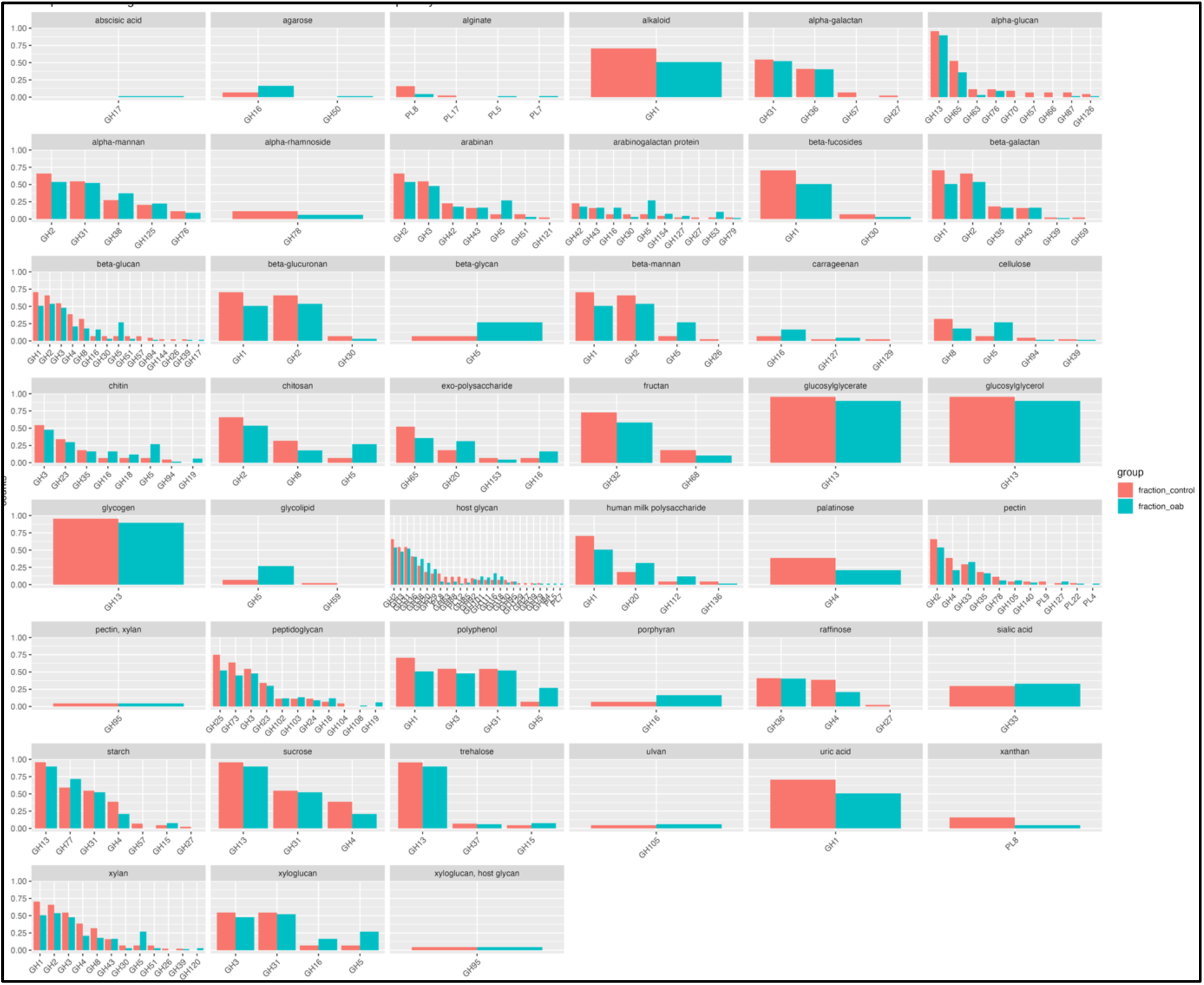
High level substrates analysis in Bladder Microbiome in OAB and Control.

**Supplemental Fig.2.**
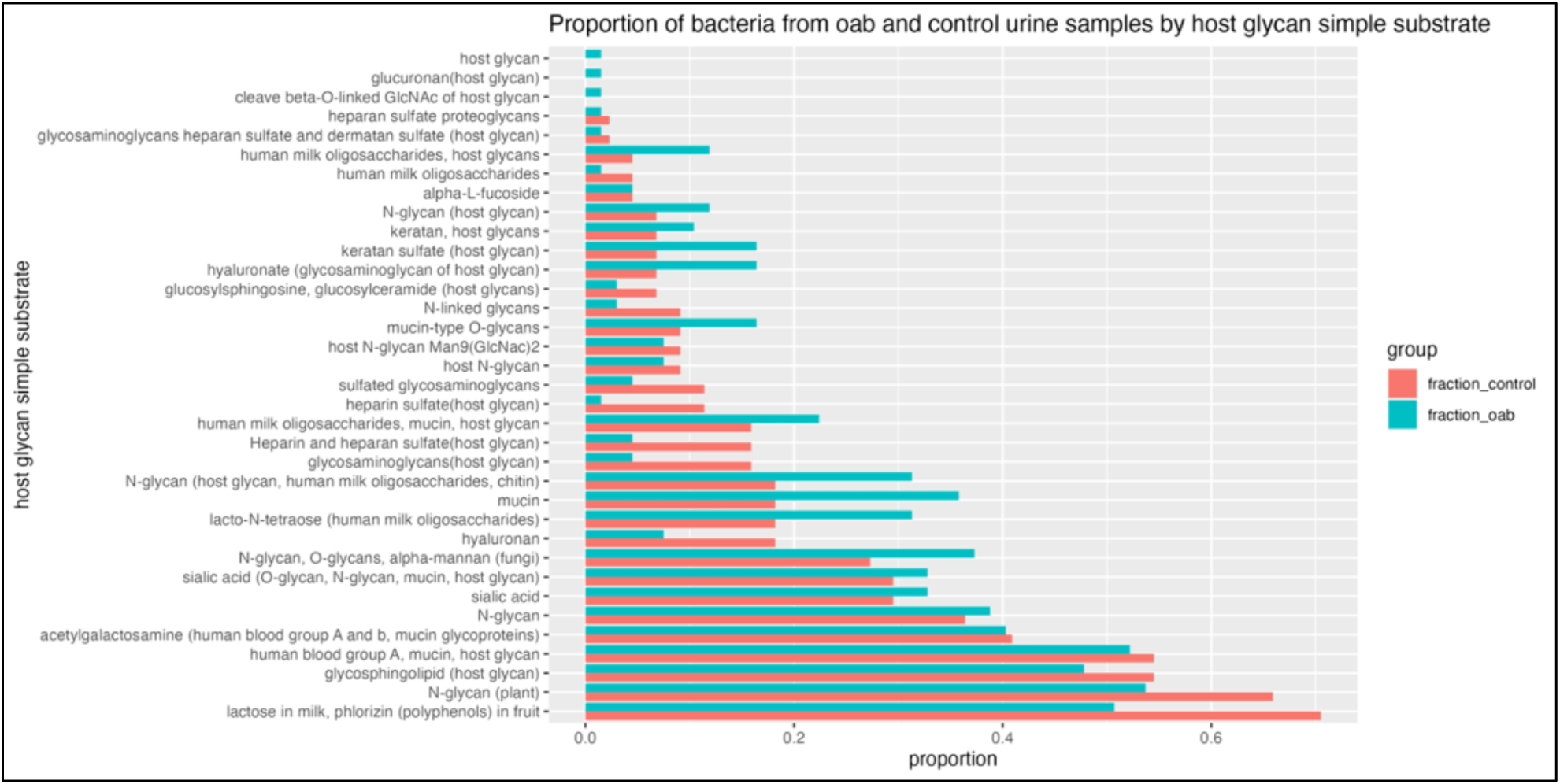
Host glycans substrates analysis in Bladder Microbiome in OAB and Control.

